# Spontaneous genomic variation as a survival strategy of nosocomial *S. haemolyticus*

**DOI:** 10.1101/2022.05.17.492267

**Authors:** Ons Bouchami, Miguel Machado, João André Carriço, José Melo-Cristino, Herminia de Lencastre, Maria Miragaia

## Abstract

*Staphylococcus haemolyticus* is one of the most important nosocomial human pathogens frequently isolated in bloodstream and medical devices related infections. This species is notorious for its multidrug resistance and genome plasticity. However, its mechanisms of evolution and adaptation are still poorly explored. In this study we aimed to characterize the strategies of genetic and phenotypic diversity in *S. haemolyticus*. Here, we analyzed an invasive *S. haemolyticus* strain, recovered from a bloodstream infection, for genetic and phenotypic stability after serial passage *in vitro* (>400 generations) in the absence and presence of sub-inhibitory concentrations of a beta-lactam antibiotic. We performed PFGE of the culture and five colonies at seven time points during stability assays were analyzed for beta-lactams susceptibility, hemolysis, mannitol fermentation and biofilm production. We compared their whole genome regarding chromosomal structure, gene content and mutations and preformed phylogenetic analysis based on core SNPs. We observed a high instability in the PFGE profiles at the different time points during serial passage *in vitro* in the absence of antibiotic. However, no variation was observed in PFGE patterns in the presence of beta-lactams. Analysis of WGS data for individual colonies collected at different time points showed the occurrence of six large-scale genomic deletions within the *oriC* environ (36 kbp-348 kbp) in the cell populations analyzed, smaller deletions in non-OriC environ region as well as non-synonymous mutations in clinically relevant genes. The regions of deletion and point mutations included genes encoding amino acid and metal transporters, resistance to environmental stress and beta-lactams, virulence, mannitol fermentation, metabolic processes and IS elements. A parallel variation was additionally detected in clinically significant phenotypic traits such as mannitol fermentation, beta-lactams resistance, hemolysis and biofilm formation. All the genetic variants analyzed were closely related in their core genome (13-292 SNPs). Our results suggest that *S. haemolyticus* populations are composed of subpopulations of genetic and phenotypic variants that might be affected in antibiotic and stress resistance, specific metabolic processes and virulence. The maintenance of subpopulations in different physiological states might be a strategy to adapt rapidly to a stress situation imposed by the host particularly in the hospital environment.

## Introduction

*Staphylococcus haemolyticus* is one of the most important nosocomial human pathogens frequently isolated in blood infections (including sepsis) related to implanted medical devices. It is easily distinguished from other coagulase-negative staphylococci (CoNS) by its multidrug resistant pattern, high numbers of antimicrobial resistance traits (Hira et al. 2013), formation of thick biofilms (Fredheim et al. 2009) and high phenotypic variation and genome plasticity (Takeuchi et al. 2005).

Genetic diversity is believed to result from the very large number (as many as 82 IS, of which 60 were intact) of insertion sequences (IS) elements (Takeuchi et al. 2005), which is a number much higher than that found in *Staphylococcus epidermidis* ATCC 12228 (18 intact IS) and *Staphylococcus aureus* Mu50 (13 intact IS). The large amount of IS elements is believed to contribute to its genome plasticity, through chromosomal rearrangements or deletions. However, the mechanisms of the evolution and adaptation of *S. haemolyticus* are still poorly understood.

Insertion sequences (IS) are transposable elements (less than 2.5 kb) that carry no genetic information except for transposases and short flanking terminal inverted repeats sequences (IR) (between 10 and 40 bp), which serve as recognition sites for the transposase. This enzyme usually excises the IS and inserts it elsewhere in the genome (conservative transposition), but occasionally the IS replicates during the transposition process (replicative transposition) (Chandler and Mahillon 2002; Schneider and Lenski 2004). By using mechanisms independent of large regions of DNA homology between the IS and target, these transposable elements are capable of repeated insertion at multiple sites within a genome. The impact of ISs in the overall genome architecture and gene expression can be very important, especially when present in multiple copies. These elements often cause gene inactivation (by direct integration into an open reading frame) and have strong polar effects, but can also lead to the activation (by providing the gene with a potent promoter) or alteration of the expression of adjacent genes (Zhang and Saier 2012). By changing the content of the genome, the IS elements might contribute to the innate ability of the bacteria to acquire drug resistance. Moreover, they can lead to complex chromosomal rearrangements that result in inversions or deletions, which can be very large and have impact on host adaptation (Watanabe et al. 2007).

The impact of IS transposition on *S. haemolyticus* chromosomal rearrangements have been only explored in strain JCSC1435 which has 56 copies (40 intact) of *IS1272* and *ISSha1*. In this strain genomic rearrangements occurred preferentially near the origin of replication and implicated the deletion/inversion of large chromosomal fragments, which had impact on antibiotic resistance and sugar metabolism (Takeuchi et al. 2005). However, it is not known how frequently this phenomenon occurs within the population, if it is restricted to the *ori* region, if it also occurs at the cell population level, which factors might induce it and what are the consequences for the bacteria fitness and survival. The *oriC* environ is a chromosomal region of staphylococci reported to be important for the evolution of staphylococcal species and it is significantly larger in *S. haemolyticus* compared to that of *S. aureus* and *S. epidermidis*. For example, this region integrates the staphylococcal cassette chromosome, conferring resistance to virtually all beta-lactams (Ito et al. 2001).
Another possible origin of genetic diversity in *S. haemolyticus* that can or not be related to IS is the high recombination rate described based on the examination of the sequence changes at MLST loci during clonal diversification (Bouchami et al. 2016). In particular, the per-allele and per-site recombination to mutation (r/m) rates reported for this species were 1:1 and 2.9:1, respectively (Bouchami et al. 2016).

The few studies available analyzing the population structure of nosocomial *S. haemolyticus* showed that they are genetically diverse by pulsed-field gel electrophoresis (PFGE); but belong to two main clonal lineages as concluded by multilocus sequence typing (MLST) (Bouchami et al. 2016; Cavanagh et al. 2012) and whole genome sequencing analysis (Pain et al. 2019).

In spite of the extremely high number of ISs in *S. haemolyticus* and the high recombination rate, the impact of transposition and recombination in genome architecture, population structure, and pathogenicity of *S. haemolyticus* was limitedly explored only. This study showed that chromosomal and phenotypic diversity in *S. haemolyticus* frequently occurs within a cell population, revealing a new mechanism of the evolution and adaptation of this species.

## Results

### *S. haemolyticus* invasive strain has a high genomic instability in the absence of environmental stress

In our previous study (Bouchami et al. 2016), we observed a high instability in SmaI PFGE macrorestriction patterns during serial passage *in vitro* in optimal growth conditions of the invasive *S. haemolyticus* strain HSM742. A culture originated from a single colony was the starting point of the stability assay that was performed along 34 days. The growth rate of HSM742 strain was 0.33 h^-1^, which corresponds to a duplication time of 2 h, indicating that after 34 days of serial growth *in vitro*, 408 generations have occurred. During this period, the SmaI PFGE patterns changed several times in two or more bands and in some occasions reverted to the original genotype. The PFGE patterns of *S. haemolyticus* strain HSM742 from days 4, 7, 10, 19 and 31 remained unchanged after passage (results shown in previous study; Supplemental Fig. S1) comparing to day 0. However, PFGE patterns obtained from days 13, 16, 22, 25 and 28 were distinguishable from those of the original (first) strain. We noticed that the banding pattern of strains isolated in days 13, 25 and 28 varied at three loci; one with a new band (approximate molecular weight 358 Kb) and two with missing bands (approximate molecular weights 432 and 74 Kb). The PFGE patterns of strains isolated in days 16 and 22 lost, each, one band (approximately 74-Kb and 108 Kb, respectively) and gained an additional band (432 Kb and 159 Kb, respectively).

The repetition of the stability assay in the exact same conditions and from the same original culture, but in the presence of oxacillin, showed completely different results. In particular, we found changes in PFGE patterns only at day 22. In this case, the strain of day 22 lost one band of, approximately, 159 Kb, and gained one band with 108 Kb. Additionally, the PFGE profile of the parental strain d0 in presence of oxacillin was different from that in absence of oxacillin. Interestingly, however, this PFGE pattern was similar to one of the PFGE pattern variants found in the stability assay performed in the absence of antibiotic (d22, variant V1 or V2; Supplemental Fig. S1). Results suggest that in the absence of an environmental stress there are diverse subpopulations of genomic variants in the same cell culture. However, in the presence of an environmental stress, like sub-inhibitory concentrations of antibiotic, or nutrients limitation, the most adapted subpopulation will be selected.

### Genomic variants of *S. haemolyticus* invasive strain have deletions within and outside *oriC* environ

To test this hypothesis and understand the mechanisms explaining the existence of subpopulations of genomic variants in the absence of antibiotic, we selected the seven time points, in which we observed an alteration in the PFGE patterns, to study in more detail (days 0, 13, 16, 22, 25 and 28). The culture from each of these days was plated in rich medium and five colonies on each plate were randomly picked for DNA extraction and analysis of whole genome. Draft genomes of the 35 colonies were reconstructed by *de novo* assembly followed by alignment, using Mauve (Darling et al. 2010) and the reference strain with a closed genome JCSC1435 (Takeuchi et al. 2005). Draft genomes were then aligned and visualized using the BLAST Ring Image Generator (BRIG). A visual inspection of the circular alignment of the genomes of HSM742 (Fig. 1A) revealed a relatively high similarity of the draft genomes with the reference genome (90-100%), suggesting that almost all the genome of the reference strain was covered by the Illumina sequencing performed for the 35 colonies. Also, alignment of JCSC1435 with the closed genome obtained for HSM472 strain in day 0 by Nanopore showed that the two strains were highly homologous (92% identity) and have similar chromosomal structures, suggesting JCSC1435 is an appropriate reference to use in the alignment.

**Figure 1.**
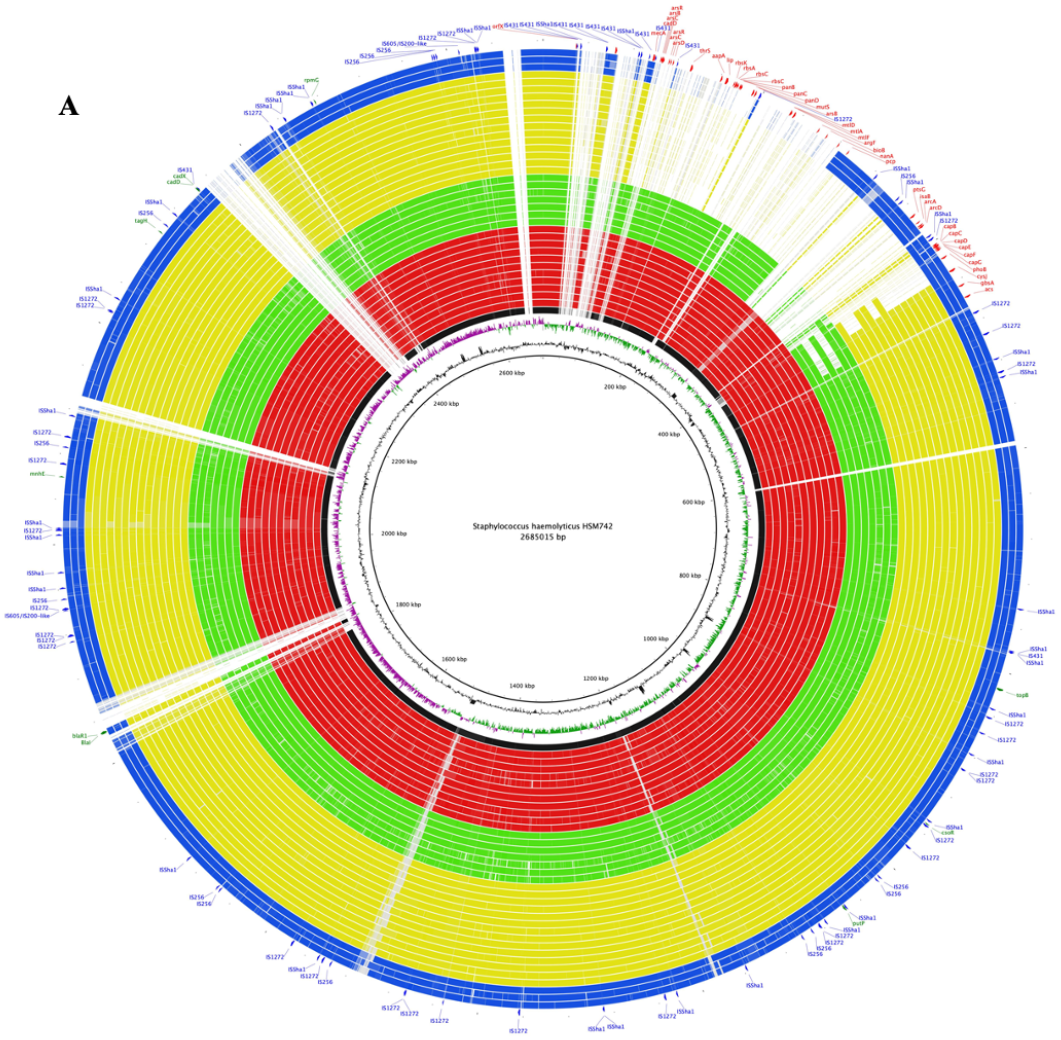
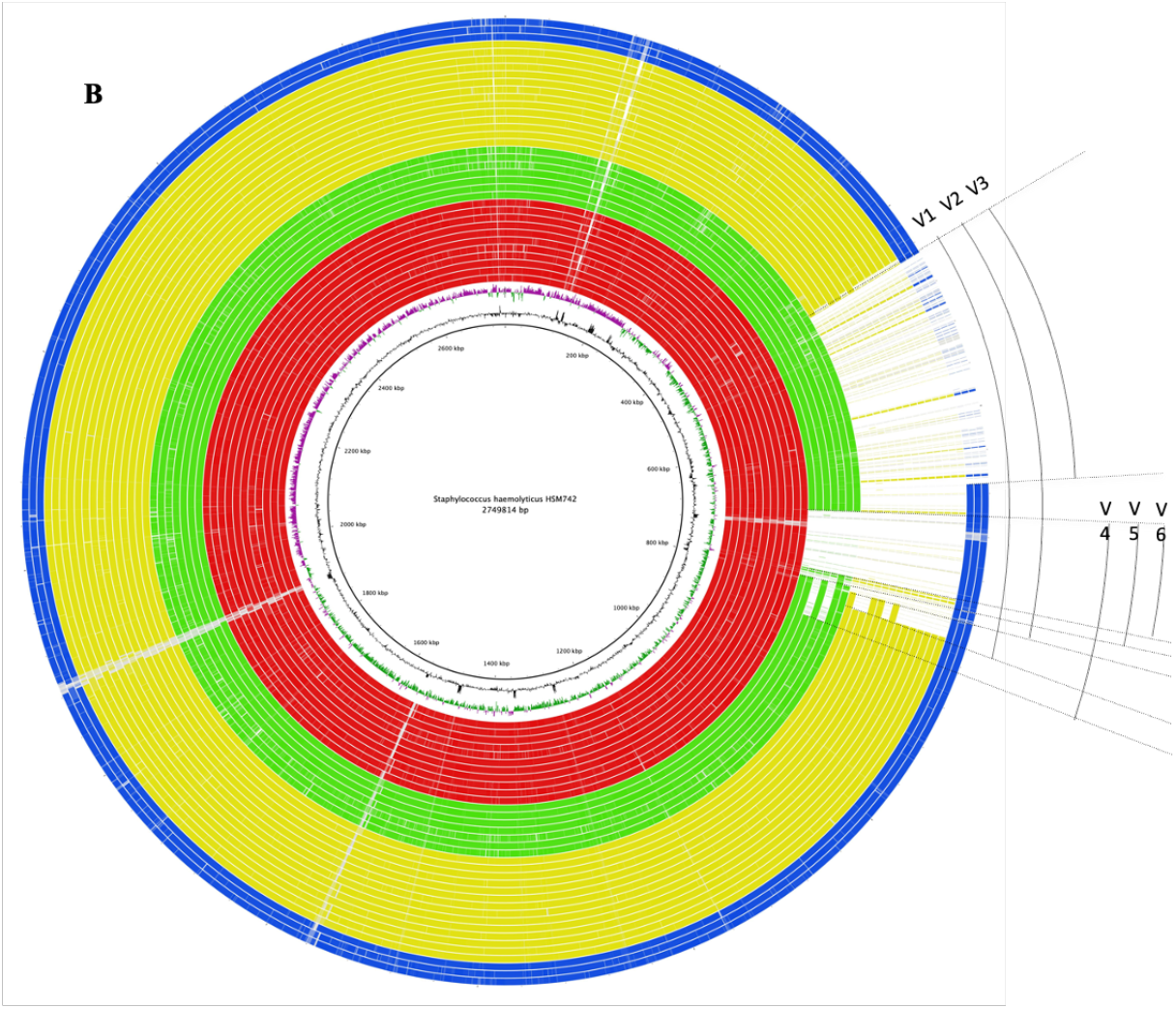
(A) Whole genome sequence analysis and comparison of JCSC1435 with other *S. haemolyticus* HSM742 strains. BRIG circular diagram of the HSM742 chromosome showing (from inner to outer), the homology based on BLASTn+ analysis of *S. haemolyticus* JCSC1435 reference genome to 35 completed HSM742 genomes (24 mannitol negative strains and 11 mannitol positive strains) (refer color-coded legend). The innermost circles represent the GC content (black), GC skew (purple/green). The outer rings show shared identity (according to BLASTn) with individual HSM742 genomes and JCSC1435 genome. BLASTn matches between 70% and 100% nucleotide identity are colored from lightest to darkest shade, respectively. Matches with less than 70% identity, or JCSC1435 regions with no BLAST matches, appear as blank spaces in each ring. Colored circles arranged from inner to outer as follow: JCSC1435 (black); Sh29/312/L2 (black); d0C1-d0C5, d13C1-C2, d16C1-C2, d28C1, d34C1 (red); d13C3 (V5), d16C3 (V5), d16C5 (V4), d25C1 (V5), d25C2 (V4), d28C2 (V6), d34C4 (V5); d13C4 (V2) (green); d13C5 (V1), d16C4 (V1), d22C1-C2, d22C3 (V1), d22C4 (V2), d22C5 (V1), d25C3-C5 (V1), d28C3-C5 (V1) (yellow); d34C2-C3 (V3), d34C5 (V3) (blue). Outer circle shows the location of the insertion sequences (blue labels and arcs) and regions of difference (deleted regions) (red labels and arcs) not present in mannitol negative strains, deleted genes outside the *oriC* (green). The image was prepared using Blast Ring Image Generator (http://sourceforge.net/projects/brig). doi:10.1371/journal.pone.0026578.g001. (B) Whole genome sequence analysis and comparison of the closed genome of HSM742d0 with other *S. haemolyticus* HSM742 strains. BRIG circular diagram of the HSM742 chromosome showing (from inner to outer), the homology based on BLASTn+ analysis of the closed genome of HSM742d0 to 35 completed HSM742 genomes (24 mannitol negative strains and 11 mannitol positive strains) (refer color-coded legend). The innermost circles represent the GC content (black), GC skew (purple/green). The outer rings show shared identity (according to BLASTn) with individual HSM742 genomes and d0C1 (sequenced by Nanopore). BLASTn matches between 70% and 100% nucleotide identity are colored from lightest to darkest shade, respectively. Matches with less than 70% identity, or d0C1 regions with no BLAST matches, appear as blank spaces in each ring. Colored circles arranged from inner to outer as follow: JCSC1435 (black); Sh29/312/L2 (black); d0C1-d0C5, d13C1-C2, d16C1-C2, d28C1, d34C1 (red); d13C3 (V5), d16C3 (V5), d16C5 (V4), d25C1 (V5), d25C2 (V4), d28C2 (V6), d34C4 (V5); d13C4 (V2) (green); d13C5 (V1), d16C4 (V1), d22C1-C2, d22C3 (V1), d22C4 (V2), d22C5 (V1), d25C3-C5 (V1), d28C3-C5 (V1) (yellow); d34C2-C3 (V3), d34C5 (V3) (blue). The image was prepared using Blast Ring Image Generator (http://sourceforge.net/projects/brig). doi:10.1371/journal.pone.0026578.g001.

The genome of the great majority of colonies (obtained from up to 34 days of serial growth and >400 generations) were highly identical in core nucleotide sequence, as shown by the small number of SNPs found (2-193 SNPs, excluding mutators) when strains were compared by core SNPs analysis (Fig. 2; Supplemental Table S1; Supplemental Fig. S2). However, six different structural genomic variants (V1-V6) were observed, when colonies’ *de novo* assembled contigs were aligned against the closed genomes of *S. haemolyticus* JCSC1435 and the HSM472-d0C1 (corresponding to a colony collected in day 1) using BRIG, or their reads mapped against the JCSC1435 strain (Fig. 1A, B). Genomic variants included deletions of different fragments sizes (313 Kb, 294 Kb, 179 Kb, 131 Kb, 82 Kb and 74 Kb), all located in the *oriC* environ right to the origin of replication between nucleotide 36000 bp and 349000 bp. Five colonies did not suffer any deletion in the *oriC* when compared to HSM742 d0C1.

**Figure 2.**
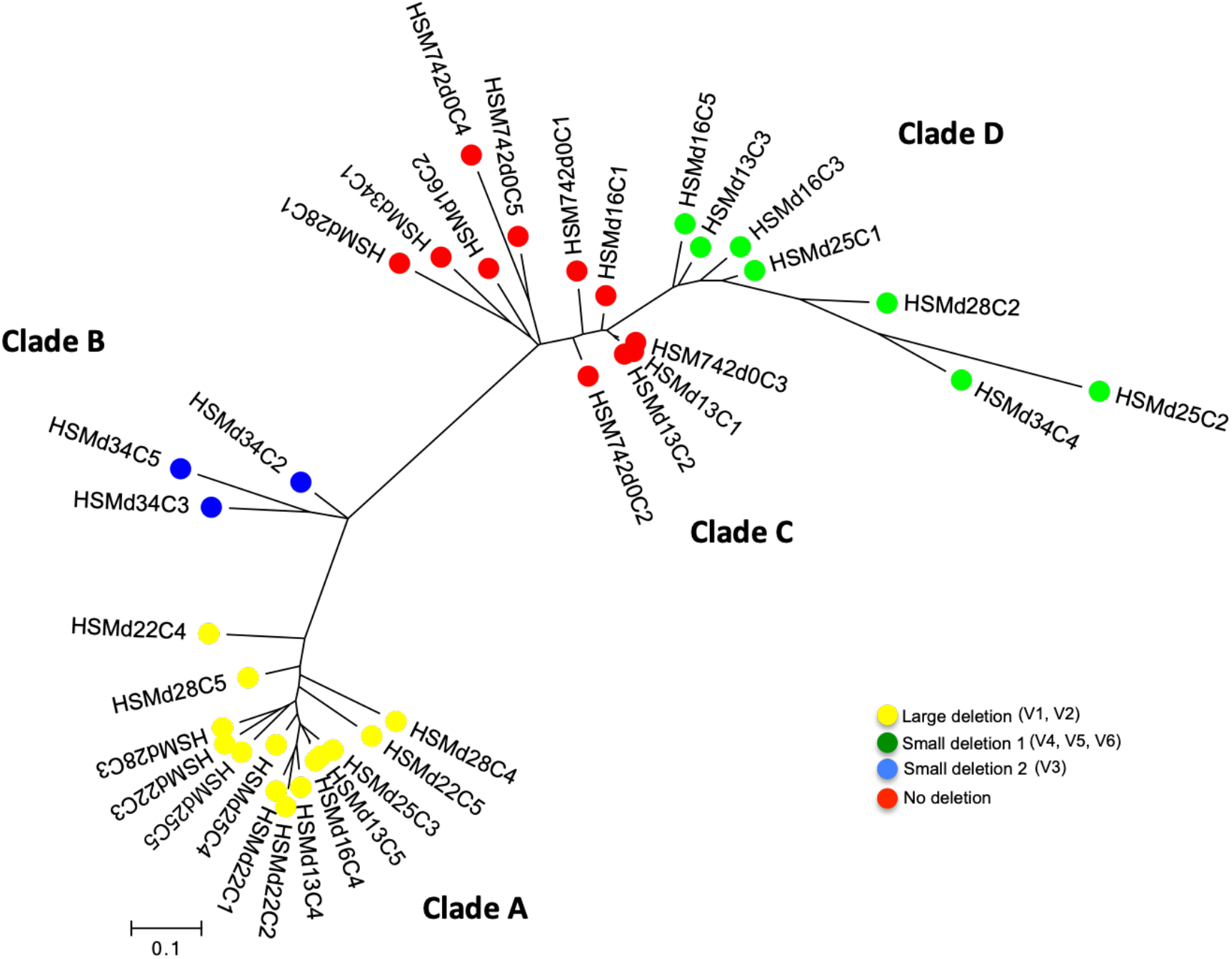
Phylogenetic RaxML tree based on 5,551 core SNPs excluding recombination, found among 35 colonies obtained from different time points in the stability assay of *S. haemolyticus* strain HSM472, using as reference the genome of JCSC1435 strain. Each node represents a strain (1 colony); nodes with identical color belong to the same cluster.

To understand which genes were contained within these regions, we constructed the pangenome of the 35 colonies and the presence/absence of genes was recorded using ROARY software. The largest deletion of 313 Kb, accounted for the loss of as many as 310 genes (Supplemental Table S2) (V1), when compared to d0C1. This deletion event was observed in 10 colonies of the stability assays in (d13C5; d16C4; d22C3; d22C5; d25C3-C5; d28C3-C5) and occurred between MaoC domain-containing protein dehydratase (36Kb) and Phosphoadenosine phosphosulfate reductase (CysH) (349 Kb). Another large fragment deletion of a similar size (294 Kb) and spanning the same chromosomal region, with the exception of the last 23bp, was also observed (V2) in four colonies (d13C4; d22C1-C2; d22C4).

There were no insertion sequences (IS) in the vicinity of the extremities of these fragments, however IS*1272*, *IS431* and *ISSha2* were part of the deletion fragment in all the 14 colonies (Supplemental Table S2). We also identified smaller deletions of 179 Kb (V3) within the same chromosomal region in three colonies (d34C2-C3; d34C5) wherein 186 genes were lost in the *oriC* environ. The upstream deletion point coincided with that of variants V1 and V2, but the downstream deletion point in these colonies was in the end of gene ABC transporter ATP-binding protein (215 Kb), instead.

The remaining three variants (V4, V5, and V6) corresponded to minor fragment deletions sizes (of 131, 82 and 74 Kb, respectively) and were detected in seven colonies (V4: d16C5, d25C2; V5: d13C3, d16C3, d25C1, d34C4; V6: d28C2). In these variants between 71, 78 and 123 genes were deleted when comparing to d0C1, and IS*1272* was located in the upstream extremity of the deletion region in 5/7 isolates. These very small deletions occurred between the MFS family major facilitator transporter, chloramphenicol:cation symporter (located 244 Kb downstrean the *oriC)* and a hypothetical protein (at 375 Kb, V4), a capsular polysaccharide biosynthesis protein Cap5G (located at 326 Kb, V5), or a FMN-dependent NADH-azoreductase, AzoR protein (at 318 Kb, V6).

We identified yet another source of variation, outside the *oriC* environ in different regions of the chromosome, wherein small deletions were also observed (Supplemental Table S3). In the 35 colonies analyzed between 71 and 313 genes were deleted outside the *oriC*, which corresponds to 16% of the total number of genes initially present in d0C1 (predicted number 1893). We could not observe a direct correlation between the gene deletions occurring outside the *oriC* environ and those occurring within the *oriC* envion, suggesting the two events are independent.

### Genomic variants of *S. haemolyticus* invasive strain are highly related

To confirm the relatedness of the 35 colonies analyzed we performed a SNPs analysis of the draft genomes obtained for each strain. The percentage of the reference genome JCSC1435 that is covered by all isolates was 65.98% implying that 1 771 593 positions from the reference were found in all analyzed genomes. A total of 5551 qualified core SNPs without recombination events was used to construct a ML phylogenetic tree (Fig. 2).

The great majority of 35 colonies differed between them in a small number of core SNPs (1-295 SNPs) (Fig. 2) (estimated medium short term mutation rate of 3.7^10^-4^ substitutions per site per year) when compared with the number of SNPs obtained when each colony was aligned to a completely distinct *S. haemolyticus* strain (JCSC1435, >5300 SNPs). The only exceptions were d0C5 and d25C2 that were more distantly related to the remaining colonies (370-1127 SNPs; 995-1127 SNPs, respectively) and might correspond to mutator genotypes. A detailed analysis of the SNPs of d25C2 (V4) and d0C5 (no deletion) confirmed the existence of several non-synonymous substitutions, when compared to other colonies, in genes involved in DNA repair function and previously associated to mutator phenotypes (Prunier and Leclercq 2005; Sinha et al. 2020). In d25C2 colony these included mutations in *mutL* (1 SNP difference with 32 variants and several SNPs with 2 variants), *recD* (1-4 SNPs with all the variants), and *recN* (1 SNP with d0C4 only). Moreover, in d0C5 colony, mutations in *mutL* (several SNPs with d34C1-C3-C4), *recO* (several SNPs with d34C4 only) and *recU* (1 SNP with d28C2 only) were additionally found.

The phylogenetic reconstruction based on core SNPs analysis grouped the 35 *S. haemolyticus* strains into four clades (A-D) (Fig. 2). The distribution of the six genomic variants coincided exactly with the phylogenetic distribution. In particular, we observed that all the colonies with large-scale deletions, including V1 and V2 variants, were grouped in the same genomic cluster A (1-52 SNPs); colonies with small deletions, including variant V3 was grouped in clade B (61-167 SNPs); and strains including the variants V4-V6 were grouped in clade D (1-144 SNPs)]. While strains without deletions were clustered together in clade C (2-193 SNPs).

We also noticed that each cluster of related genomes included colonies isolated in different days of the stability assay. For example, in cluster A we found highly related colonies isolated in days 13, 16, 22, 25 and 28; and in cluster C we found highly related colonies isolated in days 0, 13, 16, 28, and 34. On the other hand, we also observed that at day0, all the colonies tested had no deletions, belonging all to clade C, while in day 13, three colonies were deletion variants, belonging to clades A, B, and D. The proportion of the different genomic variants actually seems to vary along time. While after just a few generations (day 0) the non-deleted genome appears to predominate, in day 22 (275 generations) the large deletion variants (V1 and V2) prevailed. This change in proportion was further confirmed by testing a higher number of colonies (n=20) from each day of culture, using as a surrogate marker of the deletion event the presence/absence of mannitol fermentation (Supplemental Fig. S3; see results below). This is based on the fact that large deletions include the loss of the mannitol fermentation operon, observed by annotation of the deleted fragments. Another observation that sustains this finding is the fact that in some PFGE patterns faint bands were detected in certain time points that then become stronger in a later time point and vice versa (Supplementary Fig. S1).

Altogether, results suggest that deletion variants were already present in the starting culture (d0), or emerged in early generations, although in a low prevalence. Deletion events should have occurred a limited number of times in the population and were then maintained in the subsequent generations, evolving afterwards mainly through mutations. However, the proportion of the different deletion variants appears to have varied overtime (Fig. 2).

### Estimation of the proportion of deletion variants in the population

To establish the proportions of the variants at each time point, we took advantage of the fact that colony-variants suffering large deletions (V1, V2 and V3) had lost the mannitol operon, missing the ability to ferment mannitol, while colonies without deletions (ND) or with small deletions (V4, V5 and V6), were still able to metabolize this sugar. Actually, a total association could be observed for the five colonies tested at each time point regarding the type of variant, the presence/absence of mannitol operon (*mtlA, mtlD* and *mtlF* genes) and the corresponding ability to ferment mannitol. To increase the number of colonies analyzed and distinguish between these two types of variants at each time point, the daily cultures analyzed in this study were plated and 20 colonies picked and tested for mannitol fermentation in microtiter plates (Supplemental Fig. S3). We found that the proportion of mannitol fermenters and non-fermenters varied over time. We found that among the 20 colonies tested the proportion of mannitol negative strains was the following: day 0, 0/20; day 13 1/20; day 16, 1/20; day 22, 15/20; day 25, 18/20; day 28, 13/20; and day 34, 17/20.

### Biological functions lost through deletion events in genomic variants

As summarized in Figure 3 and in Supplemental Table S2, up to 310 genes were deleted from the largest deletion (V1) and all the other deletion variants suffered smaller deletions within this same region (V2, V3-V6) or in a region 131 Kb upstream (V4). Among the deleted genes a high proportion of genes (n= 238) encoded for hypothetical proteins. A function could be attributed for 72 genes only. These included for example genes encoding: metal transport systems and metal binding (arsenic, copper, chromium and cadmium) (*arsB, copA, chrA* and *cadD*); transcriptional regulators (*arsR, asnC, cynR, lysR*); amino acid transporters (D-serine/D-alanine/glycine transporter *aapA* gene); virulence-related genes (*isaB, lip, capABCDEFG, splE*, cell wall anchored surface proteins); sugar transport and metabolism, namely for mannitol (*mtlA, mtlD* and *mtlF)*, glucose/maltose/N-acetylglucosamine (*ptsG)*, and ribose (*rbsABCK*);metabolism of carbohydrates (*nanA*); mismatch repair (*mutS*); and restriction/modification systems (Modification methylase DpnIIA: *DpnM* gene; NgoFVII family restriction endonuclease).

**Figure 3.**
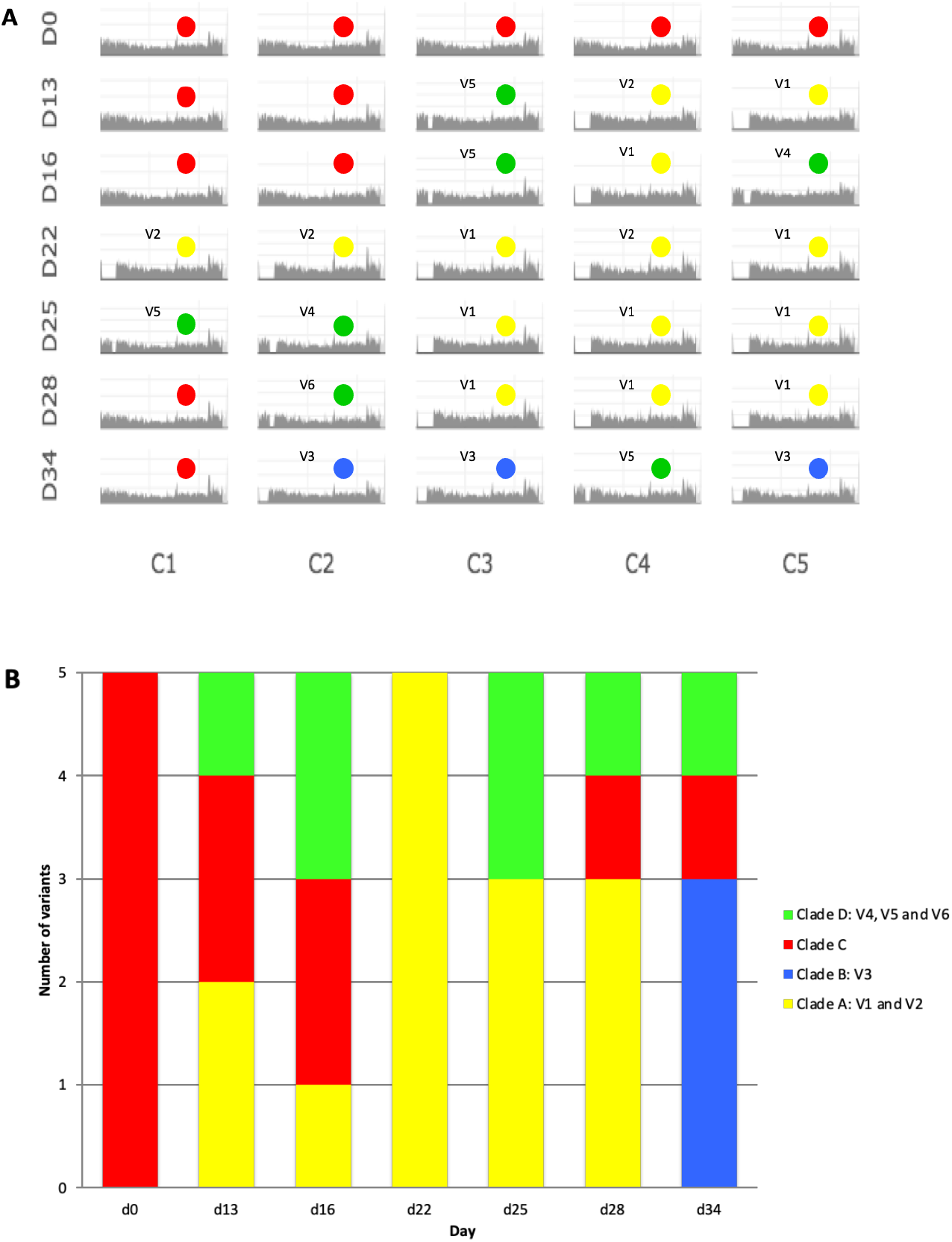
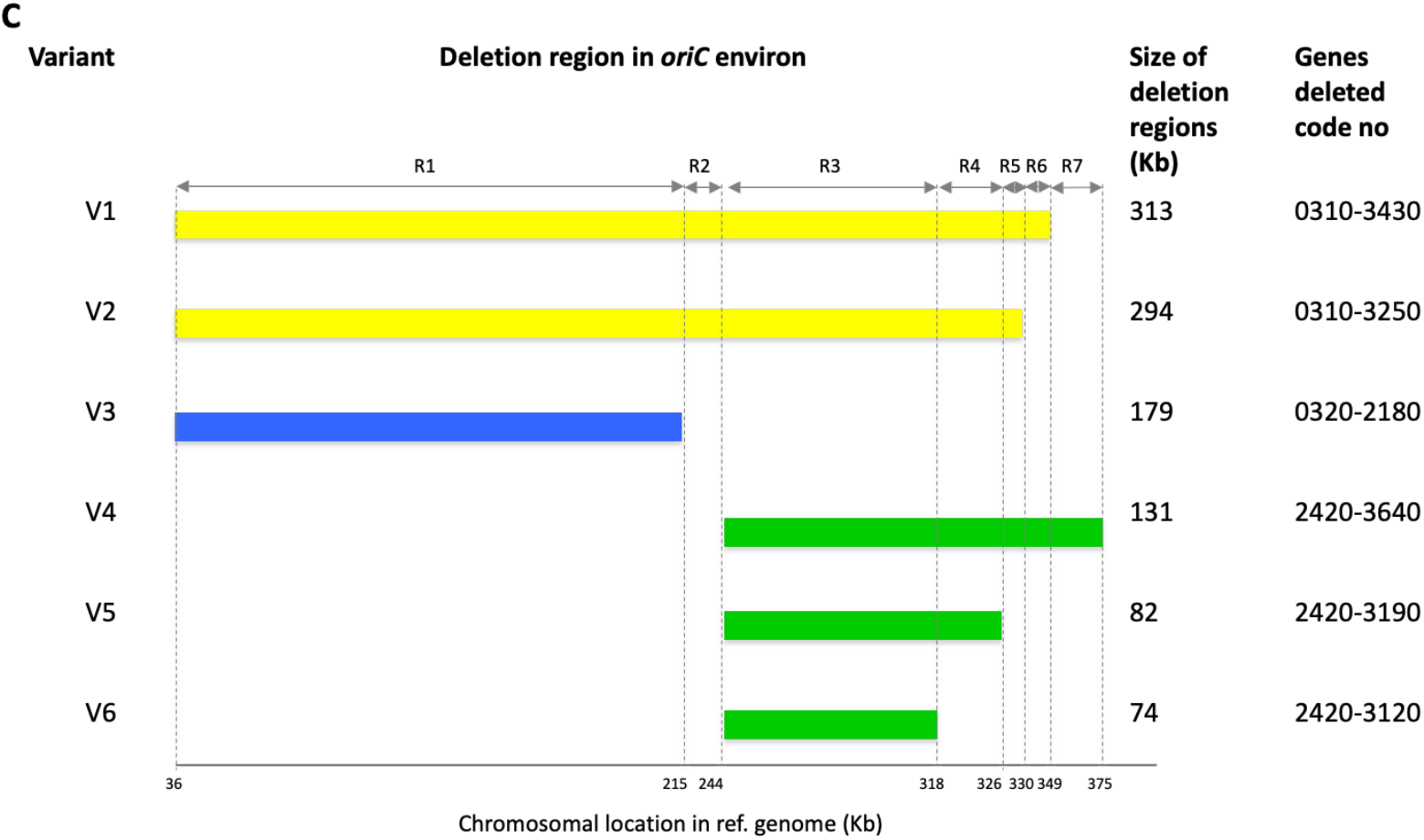
(A) Reads mapping of strain HSM742 against the reference strain *Staphylococcus haemolyticus* Sh29/312/L2. In xx axis nucleotides are in the order of the nucleotides in the reference starting from the *oriC*. In the *yy* axis is the coverage for each nucleotide position. D indicates days of passage *in vitro*. C indicates colonies. V1-V6 indicate the deletion variants and colors represent the phylogenetic clades to which the colonies belong to according to SNPs analysis; clade A – yellow; clade B – blue; clade C – red; clade D – green. (B) Distribution of deletion variants of strain HSM742 in seven time points of the stability assay. Different colors indicate the clades to which the variants belong to, according to the phylogenetic tree constructed based on the SNPs analysis. (C) Chromosomal localization and distribution of deleted genomic regions in variants of strain. Color coding has been used for visualizing the clades to which the variants belong to, according to SNPs analysis. Size of deletions are shown for each variant in kilobase (kb).

Among the 35 colonies, besides the occurrence of the large deletion in the *oriC* environ, we observed the deletion of as many as 122 additional genes outside this region, among the 35 colonies. A total of 276 genes encoding for hypothetical proteins was detected. The 122 genes with known functions included genes coding for or involved in: insertion sequences including IS*1272*, IS*431* and IS*Sau3*;,phages (prophages); metal and peptide transport (*tagH, potB, oppD, fmt, mnhE, vraG, copA*); metal binding (*csoR, sprT*); translation machinery (*rplU, rpml*, transfer RNAs); DNA replication (*dnaE, repC*);virulence, growth and survival (T-box, *rli60* and *putP*); methicillin and penicillin resistance (*blaR1, blaI*);quorum sensing and regulation of virulence (*agrC*); biofilm production (*traP, fnb, cadX, cadD, rli60*);response to osmotic and oxidative stress and acidic pH (ncRNA RsaA, RsaH) and cold shock (ncRNA RsaD); cell viability (*sipB)*.

The different deletion variants lost different functions when compared to the genome of non-deleted colonies (d0C1). For example, V3 colonies lost the ribose transporter and V4, V5 and V6 colonies lost the immunodominant antigen B (*isaB)*. However, the maintenance of several different deletion variants in the same cell population (Fig. 3A, B; Supplemental Table S2), guarantees that the entire gene pool is almost always present in the population.

### Phenotypic variation was observed during deletion events in genomic variants

To evaluate the impact of the genomic deletion events in clinically relevant phenotypes, the same 35 colonies that were used to extract DNA for WGS were used to test for mannitol fermentation, oxacillin and cefoxitin resistance level, biofilm formation and hemolysis.

Different oxacillin and cefoxitin MICs were found from colony to colony, varying from 32-256 μg/ml and 12 to >256 μg/ml, respectively. Additionally, we observed the existence of both mannitol fermenting and non-fermenting, hemolytic and non-hemolytic and biofilm producer and non-producer among the 35 colonies. Moreover, a difference in phenotypes was observed between colonies isolated from the same time point, namely in cefoxitin MICs, mannitol fermentation, hemolysis and biofilm production (Table 1). This is the case of colonies collected in days 13, 16, 25, 28 and 34 that yielded both mannitol-positive and mannitol-negative results. On the other hand, the same phenotypes were found in different days of growth. For example, the cefoxitin MIC of 64 μg/ml was observed in 10 colonies collected in different days including d13C3, d13C4, d16C4, d16C5, d25C4, d25C5, d28C3-5 and d34C4.

**Table 1.**
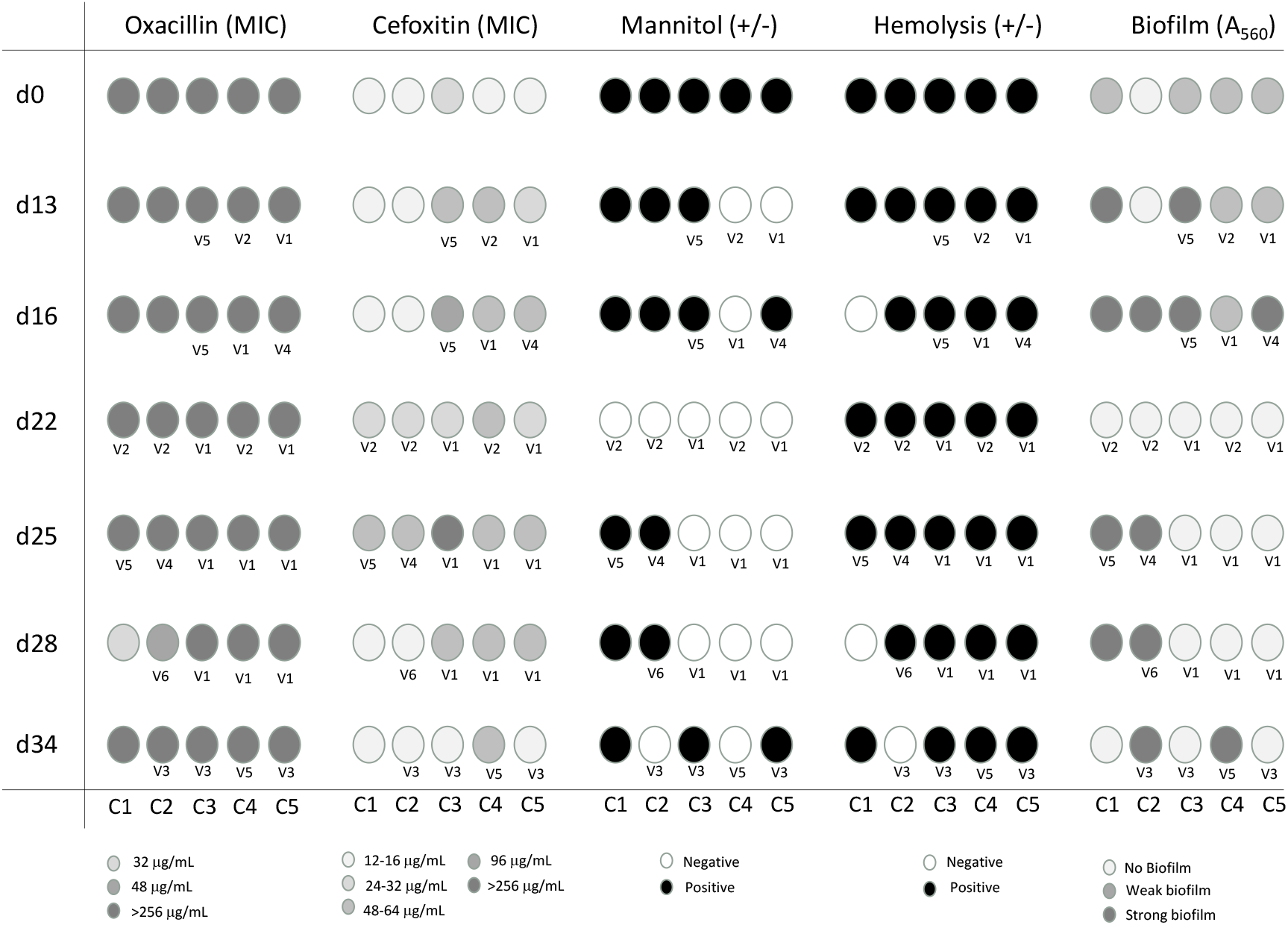
Colony to colony phenotypic variation of *S. haemolyticus* strain HSM72 after 34 days of serial growth *in vitro*. Alterations in phenotypes are shown as a gray scale

### Establishment of association between phenotypic and genotypic changes

In order to identify potential links between phenotypic changes observed and the deletions events occurring in the *oriC* environ, described in the different colonies, we used a targeted approach in which we compared phenotypes obtained for each feature tested with the type of deletion variants found for each colony (V1-V6) and looked for genes potentially associated to the phenotype within the deletion region. Additionally, to understand if genes deleted in the region outside the *oriC* environ could be implicated in the emergence of phenotypic variants, we performed an untargeted association analysis. For this purpose, the pangenome of the 35 colonies was constructed using ROARY software and the presence/absence of genes was analyzed. Besides, we also searched for the occurrence of non-synonymous SNPs in genes potentially associated to the phenotypes tested.

All deletion variants, except V3 and V6, were associated to an increase in cefoxitin MIC, when compared to the colonies without deletions. The results suggest that genes included in the region of deletion that is common to V1, V2, V4, V5 but that is present in V3, V6 and non-deleted colonies (regions R2 and R4 in Figure 3C), should be important for this change in phenotype. Among all genes contained within R2 and R4 regions, those whose functions have been previously associated to beta-lactam resistance regards to genes (*opuD*) encoding transport systems for osmolytes, such as choline/carnithine/betaine. These mechanisms were previously described to be upregulated and associated to increased susceptibility to beta-lactams (decrease in MIC) in *femAB* mutants in *S. aureus* and were found to be upregulated in methicillin resistant *S. aureus* (MRSA) heterogenous phenotype when compared to a homogenous phenotype (Keaton et al. 2013; Beenken et al. 2004; Hübscher et al. 2007).

We also observed that the variant V6 had an oxacillin MIC=48 ug/mL, while all the other deletion variants as well as non-deleted colony had a MIC>256 ug/mL, but in this case no region of deletion is specific of V6 only, and no genes were only found present in this variant only, as deducted from the presence/absence analysis. Analysis of SNPs of the variant V6 (d28C2) showed mutations in *recU* gene (1 SNP difference with all the variants; non-synonymous substitution) and in *femB* gene (1 SNP difference with d0C5 and d25C2 (V4); nonsynonymous substitution) suggesting that the phenotype alteration could be associated to mutations in these genes, described previously to be involved in oxacillin resistance (Table 1).

There was also an almost perfect correlation between the loss of mannitol fermentation ability and the deletion variants V1, V2 and V3. Although the deletion of the region that is common to these three variants (R1 region) contains as many as 188 genes, the loss of mannitol fermentation ability is most probably explained by the deletion of the *mlt*operon, contained within this R1 region.

Regarding biofilm phenotype the association between the phenotype and genotype appears to be more complex. While all the V1 and V2 colonies showed decreased biofilm formation when compared to the non-deleted colonies, in V4, V5 and V6 colonies biofilm production was increased, and V3 showed variable results. The genes associated to the decrease in biofilm may be within the deletion region that is specific of V1 and V2 only (R2 region in Table 2 and Figure 3C). Among the few annotated genes in this region, those encoding for oligopeptide transport were previously found to be upregulated in biofilm when compared to planktonic growth in *S. aureus* (Beenken et al. 2004; Graf et al. 2019) and might be also responsible for the decrease in biofilm when deleted in variants V1 and V2. The genes associated to an increase in biofilm formation should be associated to a deletion region that was common and specific of V4, V5 and V6 or to genes that are specifically found in these variants and not found in V1 and V2 or in the non-deleted colonies, but according to the deletion chromosomal locations, this is an impossibility. It is thus conceivable that the strong biofilm phenotype might be due to mutations occurring in regions of the chromosome different from *oriC* environ or new regulatory mechanisms induced by the deletion event. Analysis of SNPs showed mutations in *cadD* and *arsC* genes (nonsynonymous substitutions) in strong biofilms producers comparing to biofilm weak producers or non-producers suggesting that the phenotype alteration could alternatively be due to mutation in these genes that have been previously shown to be involved to biofilm formation (Gill et al. 2005; Zhou et al. 2020; Vernyik et al. 2020; Wu et al. 2015). Regarding hemolysis no clear alteration in phenotype was observed and the small variations in the hemolysis halo observed were not correlated with any particular deletion variant, gene or mutation.

**Table 2.**
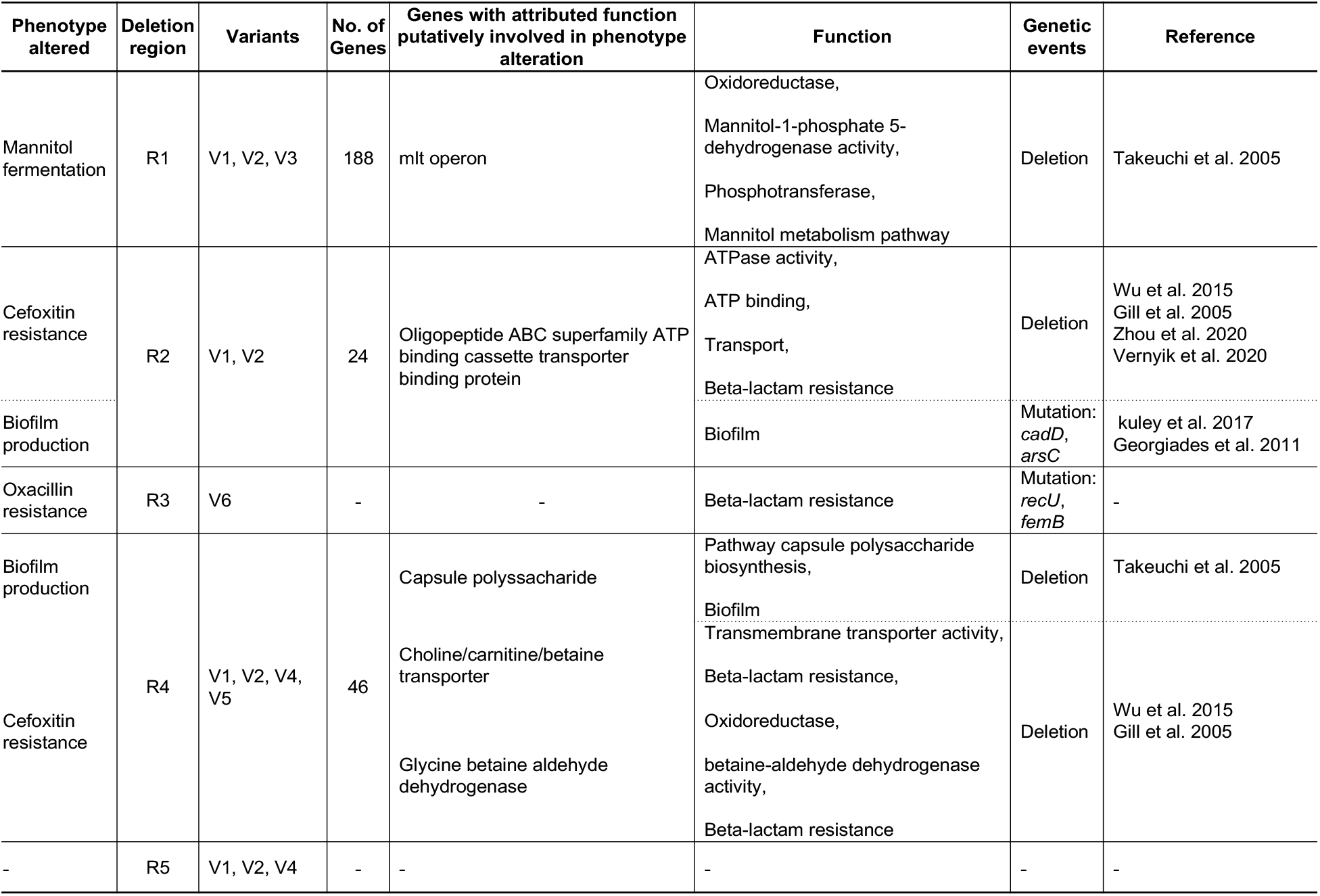
Deletion regions and genetic determinants putatively involved in phenotype alteration in *Staphylococcus haemolyticus* HSM72 strain variants

## Discussion

In the present study we have gained a deeper understanding of the mechanism of chromosomal and phenotypic diversity in *S. haemolyticus* within a cell population by analyzing a nosocomial strain belonging to the most prevalent *S. haemolyticus* clonal type. This strain was analyzed for genetic and phenotypic stability after serial passages *in vitro* in the presence and absence of a physiological stress, using whole genome sequencing, phylogenetic and pan-genome analysis.

In our previous study (Bouchami et al. 2016), we found that the SmaI PFGE macrorestriction patterns of the invasive *S. haemolyticus* strain HSM742 were highly unstable during serial growth *in vitro* in optimal growth conditions. In this study, by sequencing five individual colonies from seven time points (n=35), we found that the variability generated during serial growth in the absence of antibiotic was due to the existence of sub-populations of genomic variants deriving from the same ancestral strain. This was supported by the low number of SNPs found when the core genome of the 35 colonies was compared. Although the genomes of all colonies were highly related, we observed six different structural genomic variants (V1-V6) which included deletions of different fragments sizes, all located in the *oriC* environ right to the origin of replication. This chromosomal region was previously shown by others to be highly variable (Takeuchi et al. 2005). Moreover, colonies’ genomes suffered small deletions and nonsynonymous mutations in genes located in regions of the chromosome outside the *oriC* region. The deletions observed were most of the times associated to insertion sequences that were either within the deletion fragment or in its extremity, suggesting they might be involved in the deletion process. However, recombination, previously shown to be frequent in *S. haemolyticus* (Bouchami et al. 2016), and mutation events might be also contributing to this diversity.

When the SNPs-based phylogenetic tree of the 35 colonies was constructed we found that each of the six deletion variants were grouped in a specific cluster of the tree, which was different from the cluster containing colonies without deletions. Every cluster included colonies isolated in different days of the stability assay and the same variant was detected in different points in time. Additionally, the proportion of the different genomic variants varied over time. Altogether, these results suggest that deletion variants were created in the population a limited number of times early during serial growth and were then maintained in the subsequent generations with their proportions changing overtime. However, when serial growth was performed in the presence of sub-inhibitory concentrations of oxacillin, specific variants, namely those corresponding to a large deletion, were selected, as suggested by the analysis of the PFGE patterns along serial growth in these conditions. These results point towards the existence of subpopulations of variants as a survival strategy to counteract stress, wherein the most adapted variant will be the one that will be selected and prevail.

In these variants an impressive number of between 71 and 310 genes in the *oriC* region were deleted when compared to a colony from the starting culture (d0C1). Deleted genes included a plethora of different functions, namely those encoding carbohydrates, sugar, metal and aminoacid transport and metabolic systems and metal binding; transcriptional regulators; virulence-related genes; mismatch repair; and restriction/modification systems. We also observed the occurrence of deletion of additional genes outside the *oriC*, such as: insertion sequences (transposases) and phages (prophages); peptide transport; translation machinery; DNA replication; virulence, growth and survival; methicillin and penicillin resistance; quorum sensing; biofilm production; response to osmotic, oxidative and acidic stress and cold shock; and cell viability and hemolysis. Furthermore, when strains were compared by SNPs analysis, non-synonymous mutations in relevant genes were additionally detected. Moreover, we were able to demonstrate that the gene deletions and mutations detected were frequently paralleled by changes in clinically relevant phenotypes such as biofilm formation, beta-lactams resistance, mannitol fermentation and hemolysis.

Altogether, deletions in *oriC* region and outside *oriC* represent near a quarter of the *S. haemolyticus* chromosome and can constitute a mechanism of genome reduction by which bacteria would gain fitness (Kuley et al. 2017; Vernyik et al. 2020). On the other hand, it could be a mechanism for specialization of *S. haemolyticus* to survive in a specific niche as seen previously for other bacterial species (Georgiades and Raoult 2011; Kuley et al. 2017; Rolain et al. 2013). Although the different deletion variants lost distinctive functions, the maintenance of several different deletion variants in the same cell population, as the one observed in our study, guarantees that the entire gene library is present in the population at all time points. The existence of such a genetic diversity, would allow that the most adapted variant would emerge from the population, when faced with a new environmental challenge. This was the case when the strain under study was grown in the presence of oxacillin, wherein the most beta-lactam resistant variant (V1 and V2, MIC >256 μg/mL) was selected.

Although the study described here was totally performed *in vitro*, a recent study by Both *et al*.detected the presence of *S. epidermidis* deletion variants during prosthetic joint infections (PJI) (Both et al. 2021), suggesting that the mechanism described here for *S. haemolyticus* could actually occur *in vivo* and be a common strategy of coagulase-negative staphylococci to circumvent host immune defenses and stresses imposed by hospital environment.

Overall, our results suggest that *S. haemolyticus* populations are composed of subpopulations of genetic variants that might be affected in their growth, gene expression level, stress resistance, specific metabolic processes and virulence. The high genetic and phenotypic variability observed in the most epidemic *S. haemolyticus* clonal type appears to be the result of IS dependent and also IS independent events such as recombination and mutation events. The maintenance of subpopulations of cells in different physiological states might be a strategy to adapt rapidly to environmental stresses imposed by host or hospital environment.

## Methods

### Bacterial strains

The *S. haemolyticus* HSM742 strain used in this work was isolated from the blood culture of a 56-year-old male patient at a hospital in Portugal in 2010. The strain belonged to the most epidemic (most frequent and widely disseminated) *S. haemolyticus* clonal type (ST1, CC29) (Bouchami et al. 2016)[9]. Strains *S. haemolyticus* JCSC1435 and *S. haemolyticus* Sh29/312/L2 were used as references for WGS, the methicillin-resistant *S. aureus* (MRSA) strain WIS was used as a control for stability assays and *S. aureus* ATCC 29213 from the American Type Culture Collection (ATCC) was used as a control for antimicrobial susceptibility testing.

### Assessment of genomic stability *in vitro*

*S. haemolyticus* strain HSM742 was subjected to serial passages on tryptic soy broth (TSB). A single colony was transferred to Tryptic Soy Broth (TSB) (Difco, Detroit, USA) and grown 24 hours at 37°C. Cultures were daily transferred to fresh liquid medium (1:100 dilution) for 34 days (Bouchami et al. 2016)[9]. To assess for genomic stability, the same set of experiments was carried out under antibiotic pressure using TSB supplemented with sub-inhibitory concentrations of oxacillin (MIC/4) (Oxoid, Basingstoke, UK). MRSA isolate WIS was used as an internal control for the stability assay. Serially grown populations were characterized by PFGE (Chung et al. 2000).

### Evaluation of phenotypic stability *in vitro*

Serially grown populations or colonies were tested for oxacillin and cefoxitin minimum inhibitory concentrations (MICs), hemolysis halo, mannitol fermentation and biofilm production.

Oxacillin and cefoxitin MICs were determined using E-tests (AB BioMérieux, Solna, Sweden) according to CLSI recommendations (CLSI 2014). Hemolysis was tested by spotting 5 μl drops of an overnight bacterial culture on the surface of blood agar plates for 48h at 37°C (Boerlin et al. 2003). Mannitol fermentation was tested by inoculation of isolates onto mannitol salt agar (Becton, Dickinson and Company, Le Pont de Claix, France) followed by incubation for 24-48h at 37°C. Biofilm formation was detected by the microtiter plate assay method (Fredheim et al. 2009; Christensen et al. 1985).

### Assessment of cell population variability

To assess variability within the same cell population, five colonies at seven time points during stability assays (n=35 colonies) were analyzed for phenotypic features and whole genome sequencing (WGS).

Serial dilutions of cultures corresponding to days 0, 13, 16, 22, 25, 28 and 34 were spread over the surface of TSA agar and incubated overnight at 37°C. Five colonies (~17-100% of the population) were selected randomly from the plates of the ancestral strain HSM742 (day 0) and from subsequent generations (days 13, 16, 22, 25, 28 and 34). The half of each colony was used for performing the phenotypic assays (oxacillin and cefoxitin resistance, hemolysis, mannitol fermentation ability and biofilm production) and the other half was used for DNA extraction for whole genome sequencing (WGS).

### Whole genome sequencing and *de novo* assembly

Genomic DNA was isolated from half a colony using the Qiagen DNeasy Blood & Tissue Kit (Qiagen, Limburg, The Netherlands) and sequenced by Illumina MiSeq system. Libraries for genome sequencing were constructed using the Nextera XT DNA sample preparation kit (IIIumina) and sequenced using 150 bp pair-end reads with an estimated coverage of 100x. After trimming, the reads were assembled *de novo* into contigs using the CLC Genomics Workbench 9.0 (Qiagen, Hilden, Germany) analysis package with default parameters.

Additionally, pure and high molecular weight DNA was extracted from overnight culture of d0C1 by a standard phenol-chloroform extraction protocol (Sambrook and David 2006) and sequenced on a SpotON flow cell vR9.4.1 using Oxford Nanopore rapid barcoding protocol (Oxford Nanopore Technologies, ONT, Oxford, UK) MinION. Hybrid assembly using ONT data was done using Unicycler (Wick et al. 2017).

### Comparative genomic analysis

All the resulting contigs from Illumina sequencing were ordered using the closed genome of *S. haemolyticus* JCSC1435 (NCBI accession number AP006716) as a reference using Mauve (http://darlinglab.org/mauve/mauve.html) (Darling et al. 2010). Automated annotation was performed using the RAST (http://rast.nmpdr.org/) and prokka (http://www.vicbioinformatics.com/software.prokka.shtml) softwares (Aziz et al. 2008; Seemann 2014) using default settings. Genomic comparisons were undertaken using a combination of Mauve (Darling et al. 2010) and Artemis (http://www.sanger.ac.uk/science/tools/artemis-comparison-tool-act) (Carver et al. 2005). All the genomes were visualized with BLAST Ring Image Generator (BRIG) (http://brig.sourceforge.net/) (Alikhan et al. 2011) using blast with 70% and 90% for lower and upper nucleotide identity thresholds, respectively using both *S. haemolyticus* JCSC1435 and HSM742-doC1 closed genomes as references. To look for presence/absence of genes in all colonies, the pangenome was constructed using Roary pipeline v3.12 (https://sanger-pathogens.github.io/Roary/) (Page et al. 2015). 2015) with default settings.

### Core-genome Single Nucleotide Polymorphisms (SNPs) analysis

SNPs were identified separately within each strain, using CSI Phylogeny-1.4 (https://cge.cbs.dtu.dk/services/CSIPhylogeny, (Kaas et al. 2014)) pipeline, available by the Centre of Genomic Epidemiology (CGE) of the Technical University of Denmark (DTU). Mapping of the *de novo* assembled contigs against the JCSC1435 reference genome (GenBank accession number AP006716) (Takeuchi et al. 2005) was carried out using BWA version 0.7.2 (Li and Durbin 2010). Single nucleotide polymorphisms (SNPs) were identified on the basis of the mpileup files generated by SAMTools v. 0.1.18 (Li et al. 2009). The criteria used for calling SNPs were as follows: a minimal relative depth at SNP positions of 10%, a minimal Z-score of 1.96, a minimal SNP quality of 30 and a minimal read mapping quality of 25. The minimum distance between SNPs was disabled and all indels were excluded. An alignment of the SNPs was then created by concatenating the SNPs based on position on reference genome.

Gubbins software was run using default settings to detect the recombinant regions based on the SNP density (Croucher et al. 2015). The polymorphic sites resulting from recombination events were first detected and filtered out and the phylogeny was reconstructed using RAxML. The filtered SNP output was transformed into SNP distance matrix using the snp-dists v 0.62 (https://github.com/tseemann/snp-dists). A maximum likelihood tree was constructed from concatenated SNPs (from the alignment) and visualized using MEGA 7 software (Kumar et al. 2016). An estimation of the short-term mutation rate was calculated considering the number of core SNPs between a colony in day 0 with a colony collected at day 34 that did not suffer deletions.

## Data access

The whole genome raw sequencing reads data of the 35 isolates analyzed in this study have been submitted to the Sequence Reads Archives under accession number PRJNA836617.

## Acknowledgments

The authors thank Dr. José Melo-Cristino of the Instituto de Microbiologia, Instituto de Medicina Molecular, Faculdade de Medicina, Universidade de Lisboa, Lisbon, Portugal for providing *S. haemolyticus* HSM742 isolate used in this study. A part of this work was presented as an oral communication at the 12^th^ International Meeting on Microbial Epidemiological Markers (IMMEM XII), Dubrovnik, Croatia, from September 18-21, 2019.

## Author Contributions

Conceived and designed the experiments: MM and OB. Performed the experiments: OB. Analyzed the data: OB MM JC MaM. Contributed reagents/materials/analysis tools: MM, HdL. Wrote the paper: OB, MM. Revised the manuscript: MM, HdL, JC. All authors reviewed and approved the manuscript.

